# Human fibrosarcoma cells selected for very-high doxorubicin resistance, acquire trabectedin and eribulin cross-resistance, remain sensitive to recombinant methioninase, and have increased c-MYC expression

**DOI:** 10.1101/2025.06.07.658455

**Authors:** Sei Morinaga, Qinghong Han, Kohei Mizuta, Byung Mo Kang, Michael Bouvet, Norio Yamamoto, Katsuhiro Hayashi, Hiroaki Kimura, Shinji Miwa, Kentaro Igarashi, Takashi Higuchi, Hiroyuki Tsuchiya, Satoru Demura, Robert M. Hoffman

**Author notes:** ***Correspondence to*:** Sei Morinaga, MD, PhD, AntiCancer Inc, 7917 Ostrow St, Suite B, San Diego, CA, 92111, U.S.A.

## Abstract

Doxorubicin is first-line chemotherapy for soft tissue sarcoma; however, the development of drug resistance limits its efficacy. The purpose of the present study was to select very-high doxorubicin-resistant (VHDR) HT1080 fibrosarcoma cells, determine cross-resistance to second-line chemotherapy drugs, determine maintenance of sensitivity to recombinant methioninase (r-METase) alone and in combination with doxorubicin, and measure the level of c-MYC expression. VHDR-HT1080 cells were generated by cultivating HT-1080 cells in a series of step-wise progressively higher concentrations of doxorubicin, ranging from 8 nM to 15 µM, an 1875-fold increase, over a five-month period. The WST-8 reagent was used to assess cell viability. Four groups of in vitro drug-sensitivity tests were conducted, which involved both parental HT1080 and VHDR-HT1080 cells: 1) doxorubicin alone; 2) rMETase alone; 3) a combination of doxorubicin and rMETase; and 4) untreated control. The cross-resistance of VHDR-HT1080 cells to eribulin, trabectedin, gemcitabine and docetaxel was determined. The c-MYC levels in HT1080 and VHDR-HT1080 cells were measured using Western blotting. Doxorubicin had an IC_50_ of 3.3 µM against HT1080 cells and 38.2 µM against VHDR-HT1080 cells an 11.6-fold increase. The rMETase IC_50_ value for HT1080 was 0.75 U/ml and 0.59 U/ml for VHDR-HT1080. rMETase sensitized VHDR-HT1080 cells to doxorubicin. VHDR-HT1080 cells were cross-resistant to trabectedin 8.9-fold and cross-resistant to eribulin 1.87-fold compared to parental HT1080 cells. c-MYC expression was 8.4 times higher in VHDR-HT1080 cells compared to HT-1080 cells. The present results suggest rMETase may be used as a future clinical strategy to overcome super-doxorubicin resistance in soft tissue sarcoma.

## Introduction

Soft tissue sarcomas (STS) are a heterogeneous group of malignant tumors that arise from mesenchymal tissues and account for approximately 1% of all adult cancers (1). Doxorubicin is a first-line chemotherapy for STS (2). The development of acquired drug resistance to doxorubicin has limited the clinical efficacy of this drug. High levels of c-MYC expression correlate with poor prognosis and increased resistance to chemotherapeutic agents in various cancer types, including STS (3, 4). Metastatic STS remains a recalcitrant disease in need of improved therapy.

Methionine addiction is a general and fundamental hallmark of cancer termed the Hoffman effect (5, 6). Methionine restriction, including recombinant methioninase (rMETase), selectively arrests cancer cells in the late-S/G_2_ phase of the cell cycle (7, 8) and has been shown to increase the efficacy of all types of chemotherapy drugs which target cells in late-S/G_2_ (9–16). Recently we have studied the rMETase sensitivity of drug-resistant sarcoma cells (16–22).

In the present study, we aimed to establish a very high doxorubicin-resistant HT1080 fibrosarcoma cell line (VHDR-HT1080) and investigate cross-resistance to other drugs, the maintenance of rMETase sensitivity and the level of c-MYC expression.

## Materials and Methods

### Cell culture

The American Type Culture Collection (Manassas, VA, USA) the provided HT1080 cell line. Cells were grown in DMEM with 10% FBS and 1 IU/ml penicillin and streptomycin.

### Reagents

Bedford Laboratories (Bedford, OH, USA) provided doxorubicin. AntiCancer Inc. (San Diego, CA, USA) produced rMETase. The process of producing rMETase has already been published (8): Briefly, the *Pseudomonas putida* methioninase gene was cloned in *E. coli*. which was fermented to produce recombinant methioninase which was purified with a new step, diethylaminoethyl-sepharose fast-flow ion-exchange column chromatography.

### Establishment of very-high doxorubicin-resistant HT1080 (VHDR-HT1080) cells

Over the course of five months, HT1080 cells were cultured in doxorubicin concentrations that increased stepwise from 8 nM to 15 µM, an 1875-fold increase.

### Drug sensitivity assay 1: IC_50_

Cell viability was assessed using the WST-8 reagent (Dojindo Laboratory, Kumamoto, Japan). HT1080 or VHDR-HT1080 cells were cultured in 96-well plates at a concentration of 3×10^3^ cells per well in DMEM (100 μl/well). After that, the plates were incubated overnight at 37°C. The cells were treated for 72 hours with either rMETase at concentrations ranging from 0.5 U/ml to 8 U/ml or doxorubicin at concentrations ranging from 1 µM to 40 µM. Each well received 10 μl of the WST-8 solution following the culture period. The plates were then incubated at 37°C for an additional hour. In a microplate reader (SUNRISE: TECAN, Mannedorf, Switzerland), the absorption of cells treated with WST-8 was measured at 450 nM. Microsoft Excel for Mac 2016 version 15.52 (Microsoft, Redmond, Washington, United States) was used to create the drug sensitivity curves. ImageJ version 1.53k (National Institutes of Health, Bethesda, MD, USA) was used to calculate the half-maximal inhibitory concentration (IC_50_) values. Each experiment was carried out twice, in triplicate.

### Combination drug sensitivity assay

96-well plates were seeded with 3×10^3^ VHDR-HT1080 cells. The cells received the following treatment after 24 hours: 1) No treatment; 2) Doxorubicin alone; 3) rMETase alone; and 4) the combination of rMETase and doxorubicin. After 72 hours, cell viability was determined using the WST-8 regent in triplicate.

### Cross-resistance assay

The same protocol as in drug sensitivity assay 1 was used. Second-line drugs for soft tissue sarcoma, eribulin (Eisai Inc., Nutley, NJ, USA) (0.5-8 nM), trabectedin (PharmaMar, Horsham, PA, USA) (1-40 nM), gemcitabine (BluePoint Laboratories, Little Island, Cork, Munster, Ireland) (4-64 nM), and docetaxel (Accord Healthcare Inc., Durham, NC, USA) (1-16 nM), were used to determine cross-resistance of VHDR-HT1080 cells. The IC_50_ values of each drug to HT1080 and VHDR-HT1080 were determined. Cross-resistance to a drug was defined as being present when there was a increase of 1.5-fold or more in the IC_50_.

### Western immunoblotting

Proteins were extracted from HT1080 and VHDR-HT1080 cells using RIPA Lysis Buffer and Extraction Buffer (Thermo Fisher Scientific, Waltham, MA, USA) and 1% Halt Protease Inhibitor Cocktail (Thermo Fisher Scientific). 10% SDS-PAGE gels were loaded with protein samples. The samples were then transferred to polyvinylidene difluoride (PVDF) membranes with a thickness of 0.45 μm (GE Healthcare, Chicago, IL, USA). Membrane blocking was done using Bullet Blocking One for Western Blotting (Nakalai Tesque, Inc., Kyoto, Japan). The anti-c-MYC antibody was obtained from Proteintech (1:2,000, #10828-1-AP, Rosemont, IL, USA) as well as β-Actin (20536-1-AP, 1:1,000). Horseradish-peroxidase-conjugated anti-rabbit IgG (1:5,000, #SA00001-2, Proteintech) were used as secondary antibodies. The western blot was scanned using the Clarity Western ECL Substrate (Bio-Rad Laboratories, Hercules, California, USA) and UVP ChemStudio (Analytik Jena, Upland, CA, USA). The experiments were conducted three times.

Statistical analyses were conducted using EZR software (Jichi Medical University, Saitama, Japan) (24). The Welch’s t-test and Tukey-Kramer analysis were employed to ascertain the correlation between the variables. Statistically significant *p*-values were defined as less than 0.05 (25).

## Results

### Establishment of very-high doxorubicin-resistant HT1080 cells

Very-highly doxorubicin-resistant cells (VHDR-HT1080) were selected from HT1080 cells by culturing them in doxorubicin, increasing stepwise from 8 nM to 15 µM, an 1875-fold increase over a period of 5 months. The HT1080 IC_50_ of doxorubicin was 3.3 µM [data from (16)], compared to 38.2 µM for VHDR-HT1080 cells. VHDR-HT1080 cells were 11.6 times more resistant to doxorubicin than the parental HT1080 cells (Figure 1).

**Figure 1:**
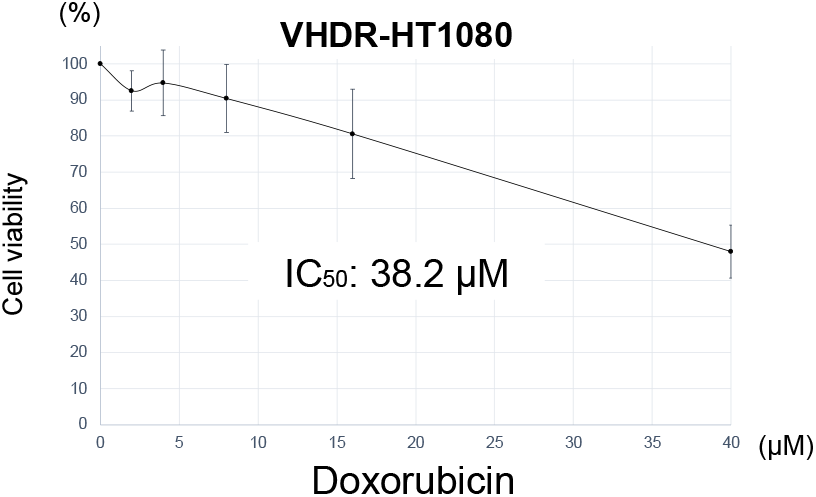
IC_50_ of doxorubicin on VHDR-HT1080 cells. Please see Materials and Methods for details. Results are shown as mean ± standard deviation.

### Determination of IC_50_ of rMETase alone on HT1080 and VHDR-HT1080

The HT1080 IC_50_ of rMETase was 0.75 U/ml [data from (10)], compared to the VHDR-HT1080 IC_50_ of 0.59 U/ml (Figure 2).

**Figure 2:**
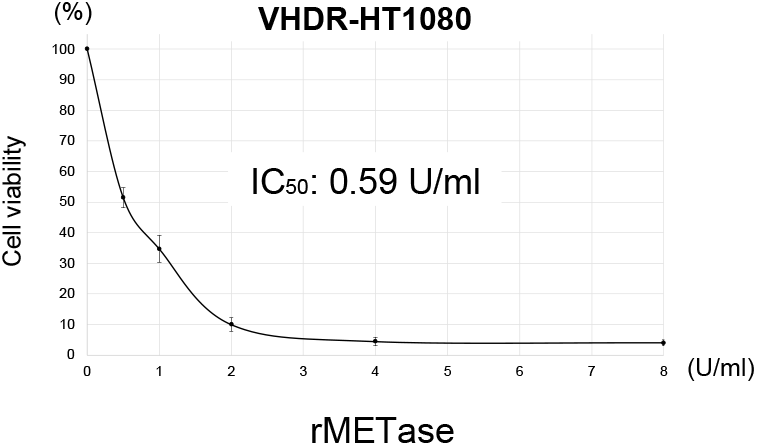
IC_50_ of rMETase on VHDR-HT1080 cells. Please see Materials and Methods for details. Results are shown as mean ± standard deviation. **VHDR-HT1080**

### Cross-resistance of VHDR-HT1080 cells to second-line STS chemotherapy drugs

The IC_50_ values for HT1080 and VHDR-HT1080 cells were 0.15 nM [data from (10)]and 0.28 nM for eribulin, respectively; 3.3 nM [data from (15)] and 29.3 nM for trabectedin, respectively; 12.8 nM [data from (25)] and 13.6 nM for gemcitabine, respectively; and 1.68 nM [data from (19)] and 1.83 nM for docetaxel, respectively (Table I).

**Table 1.** IC_50_ of HT1080 and very-highly doxorubicin-resistant HT1080 (VHDR-HT1080) cells to eribulin, trabectedin, gemcitabine or docetaxel. VHDR-HT1080 cells had cross-resistance to trabectedin and eribulin. Please see Materials and Methods for details.

|  | HT1080 (IC<sub>50</sub>) | VHDR-HT1080 (IC<sub>50</sub>) |
| --- | --- | --- |
| Eribulin (nM) | 0.15 [data from (10)] | 0.28 |
| Trabectedin (nM) | 3.3 [data from (15)] | 29.3 |
| Gemcitabine (nM) | 12.8 [data from (25)] | 13.6 |
| Docetaxel (nM) | 1.68 [data from (19)] | 1.83 |

### Efficacy of rMETase combined with doxorubicin on VHDR-HT1080 cells

The IC_50_ of rMETase for VHDR-HT1080 (0.59 U/ml) combined with the IC_50_ of doxorubicin for HT1080 (3.3 µM) inhibited VHDR cells 73.4% compared to determined control, 69.7% compared to doxorubicin alone and 15.8% compared to rMETase alone (*p*<0.05) (Figure 3).

**Figure 3:**
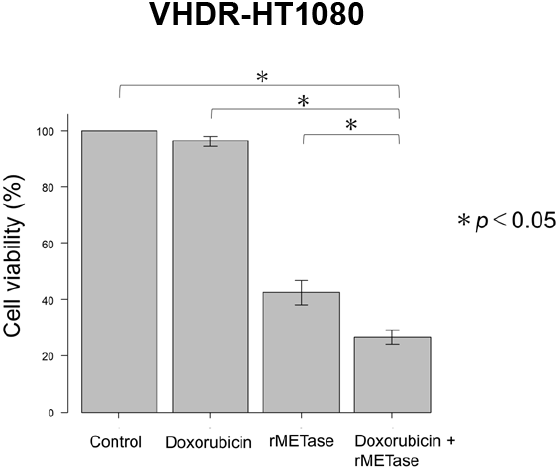
rMETase sensitized very high doxorubicin-resistant HT1080 (VHDR-HT1080) fibrosarcoma cells to doxorubicin. Control (DMEM); doxorubicin [3.3 µM (IC_50_ of HT1080)]; rMETase [0.59 U/ml (IC_50_ of VHDR-HT1080)]; doxorubicin [3.3 µM (IC_50_ of HT1080)] and rMETase [0.59 U/ml (IC_50_ of VHDR-HT1080)]. Data are shown as mean ± standard deviation. Please see Materials and Methods for details.

### Western blotting of c-MYC

c-MYC expression in VHDR-HT1080 cells increased 8.4-fold compared to HT1080 cells (*p*<0.05) (Figure 4).

**Figure 4.**
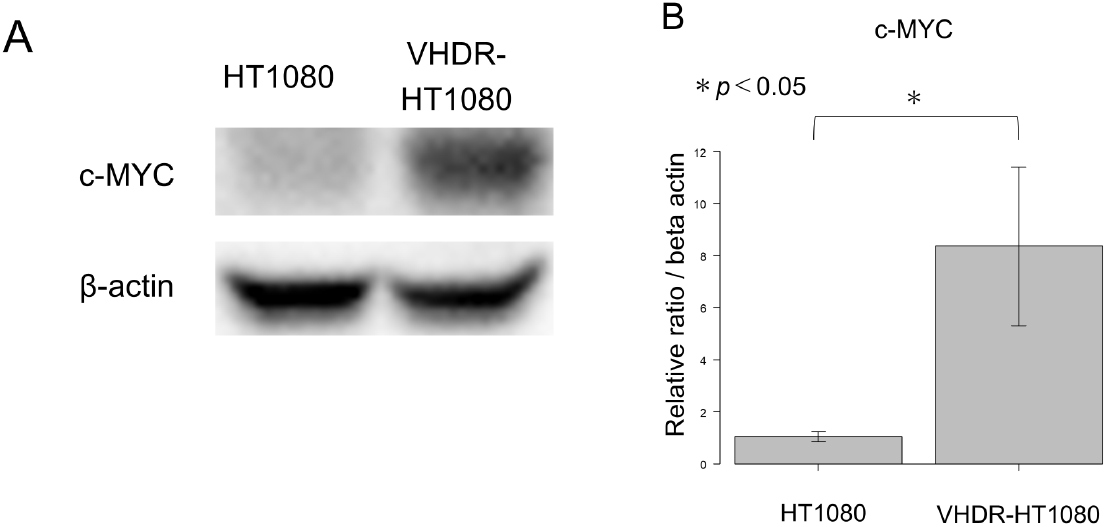
Expression of c-MYC in HT1080 and very-highly doxorubicin-resistant HT1080 (VHDR-HT1080) fibrosarcoma cells. A) Western blot of c-MYC expression in HT1080 and VHDR-HT1080 cells. B) Quantitation of c-MYC expression in HT1080 and VHDR-HT1080 cells. Data shown are representative of three different Western blots. Please see Materials and Methods for details.

## Discussion

VHDR-HT1080 was established by selecting HT1080 cells in stepwise increasing concentrations of doxorubicin (1875-fold) over five months. VHDR-HT1080 cells acquired very-high resistance to doxorubicin, with an IC_50_ value 11.6-fold greater than that of parental HT1080 cells. VHDR-HT1080 cells were cross-resistant to second-line therapy, trabectedin by 8.9-fold and to eribulin by 1.87-fold, suggesting a shared resistance mechanism.

The IC_50_ of rMETase was similar in HT1080 and VHDR-HT1080 that indicates the acquisition of very high doxorubicin resistance did not affect rMETase sensitivity. rMETase sensitized VHDR-HT1080 cells 19.8-fold to doxorubicin. These results suggest that rMETase may be effective clinically to overcome doxorubicin resistance in STS.

Further studies are needed to determine if c-MYC overexpression is linked to very high doxorubicin resistance in STS as well as the mechanism of cross-resistance to second-line STS chemotherapy in VHDR cells. The present results suggest that combining rMETase and doxorubicin may be a clinical strategy to overcome clinical doxorubicin-resistant STS that are also cross-resistant to second-line STS drugs.

## Conflicts of Interest

The Authors declare no competing interests in relation to this study.

## Authors’ Contributions

Conceptualization: SM KM RMH

Data curation: SM RMH

Formal analysis: SM KM BMK MB NY KH HK SM KI TH HT SD QH

Investigation: SM KM QH

Methodology: SM KM RMH

Project administration: RMH

Validation: KM BMK

Writing– original draft: SM

Writing– review & editing: SM, RMH.

## Acknowledgements

This article is dedicated to the memory of A.R. Moossa, MD, Sun Lee, MD, Richard W. Erbe, MD, Professor Philip Miles, Professor Gordon H. Sato, Professor Li Jiaxi, Masaki Kitajima, MD, Joseph R. Bertino, MD, Shigeo Yagi, PhD, J.A.R Mead, Ph.D., Eugene P. Frenkel, MD, Professor Lev Bergelson, Professor Sheldon Penman, Professor John R. Raper, Joseph Leighton, MD and John Mendelsohn, MD.

## Notes

### Competing Interest Statement

The authors have declared that no competing interests exist.

